# Molecular and Energetic Basis of Histidine Switch Dynamics in Respiratory Complex I

**DOI:** 10.64898/2026.01.12.698557

**Authors:** Erik Endres, Mahdi Torabi, Mai Jousmäki, Kim Vy Huynh, Cristina Pecorilla, Oleksii Zdorevskyi, Volker Zickermann, Vivek Sharma

## Abstract

The respiratory complex I in mitochondria and bacteria drives the two-electron reduction of quinone to pump protons across the membrane. The molecular basis of this catalytic reaction remains enigmatic despite significant progress in structural characterization of the complex. A highly conserved histidine residue in the distal antiporter-like subunit of its membrane domain has been shown to undergo conformational changes in molecular simulations and cryo-EM structures. However, the function of histidine switch dynamics, and energetics of its conformational transitions remain unclear. Here, by applying enhanced sampling simulations, we evaluate the energetics of the histidine switch dynamics and demonstrate that it is coupled to the charge state of lysine residues ca. 10 Å apart, which cause hydrogen bond restructuring and stabilize histidine in specific conformations. Hybrid QM/MM metadynamics simulations show that the histidine participates in gated proton transfer and may function as a proton confurcation device in complex I and related proteins.

**Importance of work:** Proton pumping by the mitochondrial respiratory chain enzymes needs to be highly directional so that the ATP production occurs as efficiently as possible during high energy demands. With the protons being hard to detect by most methods of structural biology, computer simulation approaches as applied in this work offer critical information on how molecular gates within respiratory complexes can function and help elucidate the challenging questions in the field of mitochondrial bioenergetics.

## Introduction

Mitochondrial ATP generation by ATP synthase (complex V) requires an electrochemical gradient of protons between the two sides of the inner mitochondrial membrane (Mitchell 1961). This difference in proton concentration is generated by the respiratory complexes I, III and IV, which effectively function as molecular proton pumps (Wikström, Pecorilla, and Sharma 2023), transferring protons only in one direction; that is from the negatively charged side (N-side) of the membrane to its positive side (P-side). Identifying these pumping mechanisms and how their directionality is ensured remains a challenge to this day (Kaila et al. 2008; Yang and Cui 2011). The ∼1 MDa Respiratory Complex I (RCI) is the largest complex within the mitochondrial respiratory chain and typically consists of 14 core subunits and around 31 auxiliary subunits (Parey et al. 2020). The bacterial RCIs are relatively smaller in size and consists mainly of the core subunits that form a minimal redox-driven proton pumping unit. These subunits can be divided into two domains, a solvent-exposed arm and a membrane-embedded section (Fig. 1A). NADH binds at a distal binding site and supplies electrons for the reduction of ubiquinone (UQ) at the junction of two arms (Fig. 1A). The latter triggers the pumping of protons throughout the membrane arm, generating the proton gradient for ATP synthesis. There is a general agreement on the pathway of electrons within the complex; FeS clusters, as seen in high resolution structures, clearly mark the path from the NADH oxidation site towards the UQ reduction site (Sazanov 2023; Verkhovskaya et al. 2008). The pathways for proton transfer, however, remain dubious (Fig. 1A). Some of the key questions are how substrate protons and pumped protons are channelled, how protons travel within the membrane embedded subunits, which subunits are involved in the pumping, and above all, what is the molecular basis of the redox-coupled proton pumping catalysed by RCI (Djurabekova et al. 2024).

**Figure 1.**
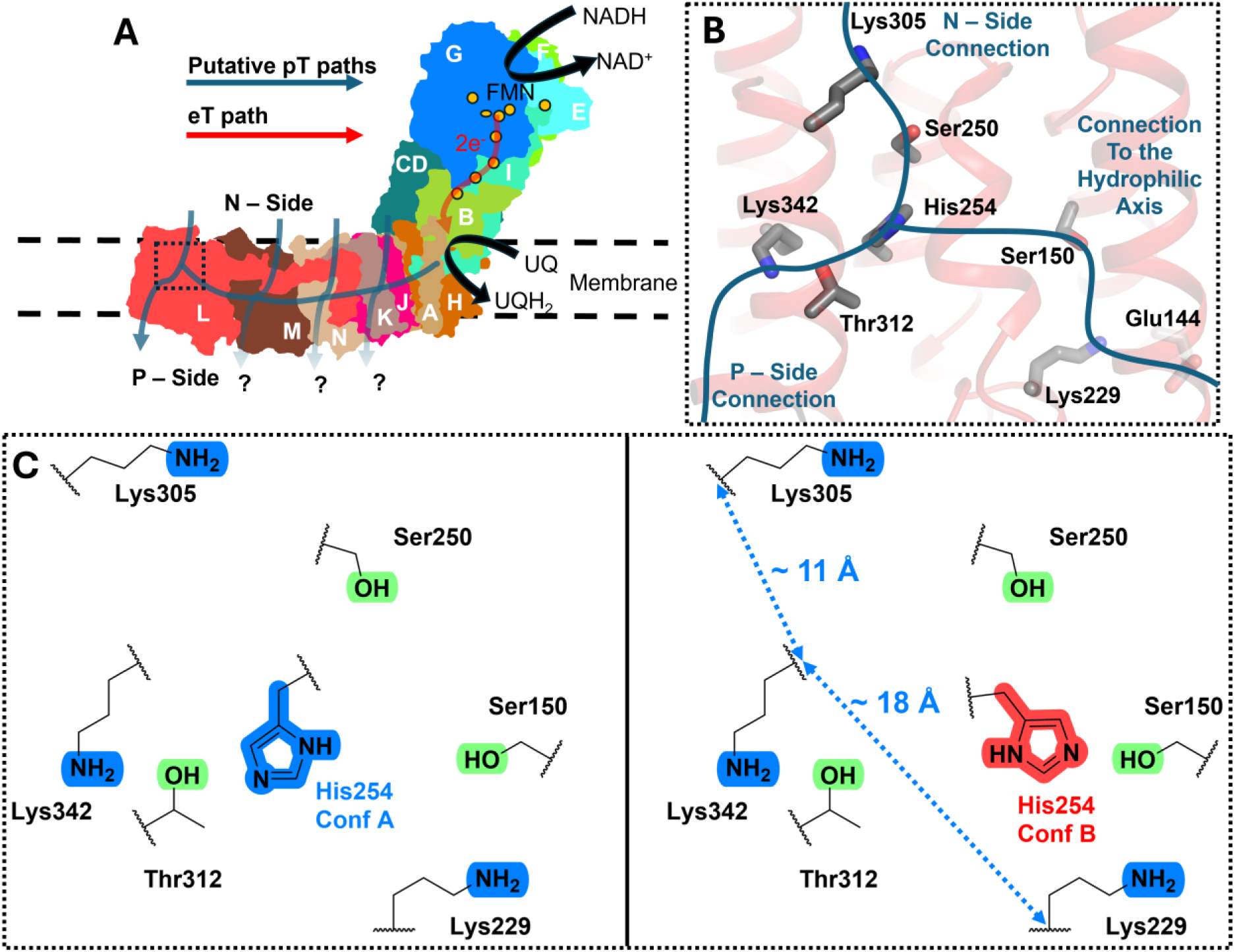
Architecture of RCI and its histidine molecular switch. (A) Bacterial RCI from *Escherichia coli,* PDB 7P7C (Kravchuk et al. 2022). Oxidation of NADH to NAD^+^ provides 2e^-^ that reduce UQ to UQH_2_. This redox reaction triggers a cascade of proton transfers, ultimately pumping protons from the negative (N) side to the positive (P) side. The electron transfer (eT) path and the putative proton transfer (pT) paths are marked in red and blue arrows, respectively. The proton pumping routes towards the P side that remain unclear are denoted with “?”. The bacterial RCI subunits are identified with single letters (e.g. L for subunit NuoL, *E. coli* RCI nomenclature. NuoCD is a fusion of the two “classical” subunits C and D). (B) His254 of NuoL connects to three putative proton transfer routes: the hydrophilic axis (right), the proton uptake site towards N side (top) and the proton release site towards P side (bottom). (C) Schematic of relevant protic residues along the putative paths (panel B) with the distances between lysine residues highlighted. His254 is shown in its two conformations (Conf A and Conf B, see text).

The three membrane-bound subunits NuoL, NuoM and NuoN (Fig. 1A) that are typically associated with proton pumping (Nakamaru-Ogiso et al. 2010; Euro et al. 2008; Amarneh and Vik 2003) are homologous to each other and share their architecture with the ancient Multiple resistance and pH adaption (Mrp) cation/proton antiporters.(Swartz et al. 2005) Among these, NuoL (or homologous MrpA in Mrp antiporter) displays several anomalous sequence features (Mathiesen and Hägerhäll 2002; Djurabekova, Haapanen, and Sharma 2020) including a strictly conserved histidine His254 (*E. coli* RCI numbering), which was found to undergo protonation-dependent conformational changes in molecular dynamics (MD) simulations of bacterial RCI (Djurabekova, Haapanen, and Sharma 2020). This histidine, which is positioned on a flexible α-helical segment, is uniquely situated at the junction of three putative proton transfer pathways, connecting to the hydrophilic axis that runs through all three antiporter-like subunits via Lys229 of a Lys-Glu pair, to a potential proton uptake path from the N-side via Lys305, and to a potential proton release path through Lys342 towards the P-side (Fig. 1B). All three lysine residues are strictly conserved, and biochemical analyses have revealed their functional significance (Nakamaru-Ogiso et al. 2010; Euro et al. 2008; Amarneh and Vik 2003) High resolution cryo-EM structures of Mrp antiporter demonstrated that histidine adopts two different conformations (Lee et al. 2022): A, pointing towards Lys342, or conformation B, pointing towards the conserved Lys-Glu-pair. In conformation A, the histidine is within a hydrogen bonding distance to Thr312, whereas in conformation B, it establishes a hydrogen bond to Ser150 (Fig. 1C). Extensive site-directed mutagenesis and computer simulations of Mrp antiporter further confirmed the functional importance of the histidine conformational dynamics in ion transport (Pecorilla et al. 2025). Building upon this data, we have proposed a model for the histidine switch dynamics in RCI (Pecorilla et al. 2025). Here, by performing atomistic MD simulations on *E. coli* RCI combined with enhanced sampling techniques, we test this proposed model in a quantitative manner by calculating the energetics of histidine dynamics in different charged and conformational states. The results posit that hydrogen bonding restructuring is central for the histidine switch function. Furthermore, metadynamics-based hybrid QM/MM free energy simulations on hydrogen-bonded pathways involving histidine provide insights on the energetics of proton transfer. We propose that the histidine switch can function as an efficient proton confurcation device in respiratory complex I and related proteins.

## Results

### Enhanced Sampling of the Histidine Switch

It is known from the modelling and simulations of Mrp antiporter that the protonation states of conserved lysine residues in the MrpA subunit (homologous to NuoL) influence the dynamics of histidine (Pecorilla et al. 2025). We therefore systematically analysed several different charged states of lysine residues in the NuoL subunit with an enhanced MD simulation sampling method (Accelerated Weight Histogram, AWH (Lindahl, Lidmar, and Hess 2014) ), using a linear combination of His254(Nδ)-Thr312(Oγ) and His254(Nε)-Ser150(Oγ) distances as reaction coordinate (RC, see Computational Methods), and obtained the energetics of the histidine switch in RCI. We find that over a broad range of protonation states, conformation A seems to be favoured over conformation B (see Figs. 2, S2 and S3). This agrees with the cryo-EM structures of *E. coli* RCI where His254 is predominantly observed to be in A conformation or in a conformation close to it. Interestingly, in several high-resolution structures of bacterial RCI, histidine is found to be stabilized in the A arrangement, in contrast to the B conformation, which is commonly observed in many mitochondrial RCI structures (see Table S2, Fig. S1). We observed that the perturbation of protonation states of selected lysine residues alters the [A]/[B] equilibrium. In contrast to ∼10 kcal/mol energy difference in favour of histidine in A conformation in state 000δ (shorthand notation describing protonation states of Lys342, Lys305, Lys229 and His254, and in that order, see also Fig. 2), protonation of Lys305 or Lys229 (state 0+0δ or 00+δ) shifts the equilibrium towards B conformation by around 2-3 kcal/mol (Figs. 2A). The protonation of Lys342 or Lys229 on top of protonated Lys305 (states ++0δ or 0++δ) also enhances the stability of B conformation. With all three lysine residues protonated (state +++δ) or in state +0+δ, the energy difference between A and B conformations reduces to only about 2-3 kcal/mol, highlighting an important role of Lys229 protonation in enhancing B population (Fig. 2). A similar lysine protonation dependent effect on histidine dynamics can be observed in its ε-nitrogen protonated tautomeric state (Fig. S2). Overall, there are only a handful of states when the energetic difference between A and B conformations is in the range of 2-3 kcal/mol.

**Figure 2.**
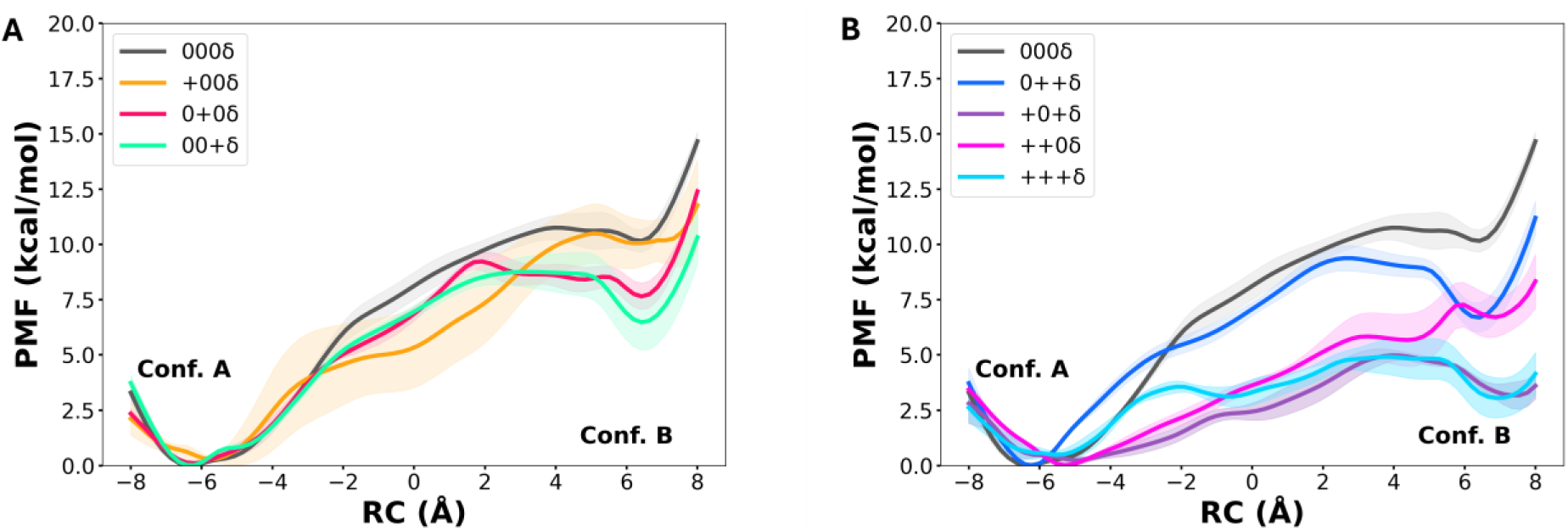
Energetics of δ-nitrogen protonated His254 sidechain conformational dynamics in different protonation states of conserved lysine residues of the NuoL subunit (Lys342, Lys305 and Lys229). The y-axis in panels (A) and (B) describe the potential of mean force (PMF, kcal/mol) with respect to the reaction coordinate (RC) on x-axis in Å (see methods). Each trace is the mean of four simulation replicas, whereas the shaded region describes the standard error of mean. The short-hand notation (+00δ) is used to describe the protonation states of lysine residues and histidine. The state +00δ corresponds to protonated Lys342, neutral Lys305, neutral Lys229 and His254 with Nδ-protonation, in that order. The Nδ and Nε tautomeric states are marked with δ and ε, respectively, whereas imidazolium histidine with p (See Figs. S2 and S3).

It is noteworthy that changing protonation states of lysine residues ca. 10-20 Å apart (Fig. 1) can have significant influence on the position of His254. To probe what stabilizes (or destabilizes) the A/B conformations, we analysed the simulation trajectories. When histidine occupies the A position (Fig. 2), the region between His254 and Ser150/Lys229 is packed with several bulky lipophilic side chains of Leu153, Ile154, Cys149, Val249, Ile254, Met258, Trp238, Leu239, Val259, Val146, including the mitochondrial disease locus Phe123 (Hoeser et al. 2025; Pecorilla et al. 2025) (Fig. 3). These essentially impart a hydrophobic barrier towards the B conformation. Only when Lys229 (and also Lys305) is modelled protonated, hydration in the region increases, leading to the destabilization of the hydrophobic blockage and establishment of the B conformation of His254 (Fig. 3).

**Figure 3.**
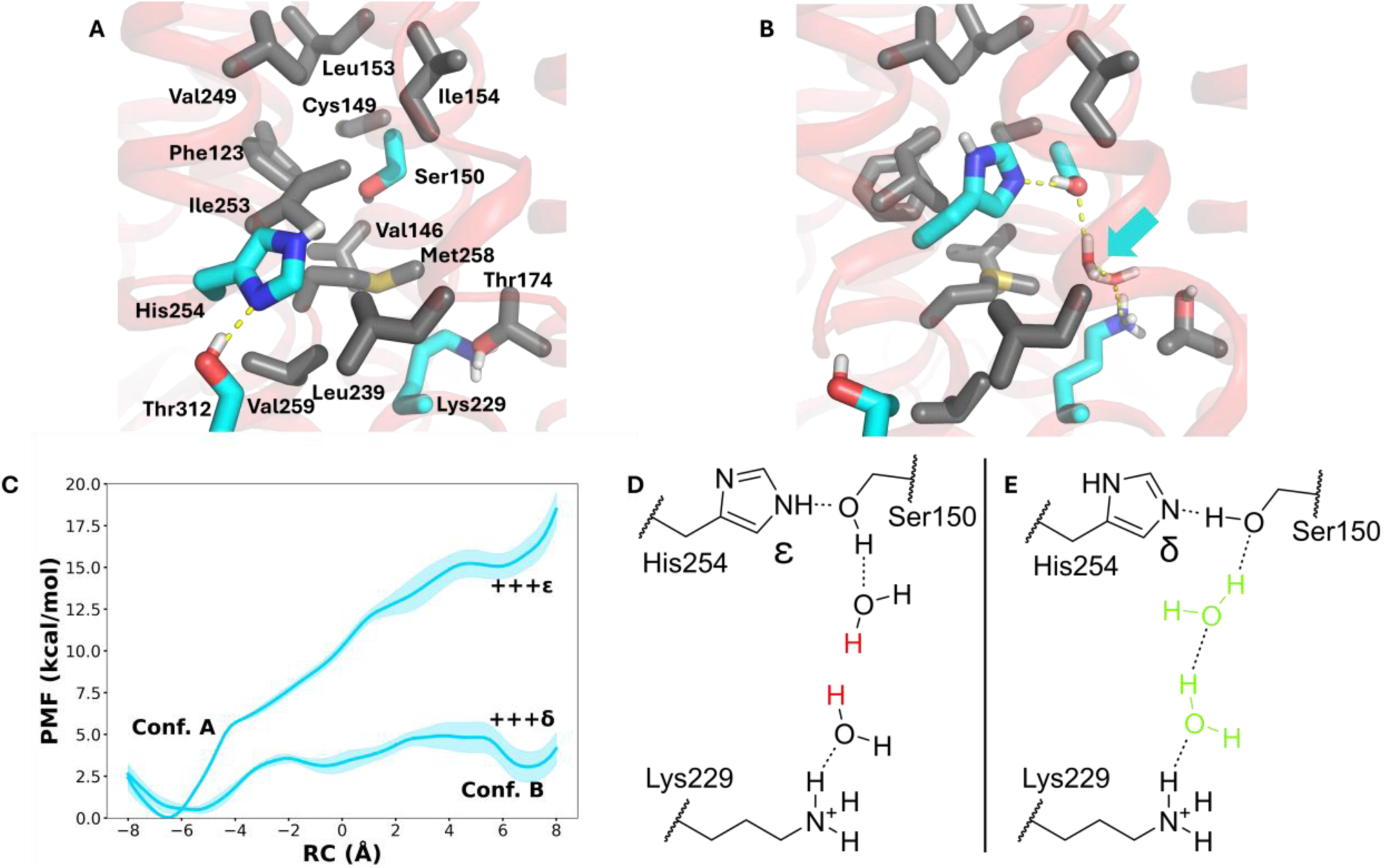
Structural changes upon histidine dynamics and its dependence on tautomeric states. The movement of His254 from A position (A) to B position (B) is blocked by a cluster of hydrophobic residues. The hydrophobic cluster rearranges upon charge-induced hydration in the region in part caused by protonation state changes of surrounding lysine residues (see text). The hydrogen bonds of His254 with Thr312 in A position and with Ser150 in B position are shown (yellow dotted lines), along with the water molecules (cyan arrow) that populate the region upon lysine protonation state changes (panel B). The snapshots shown in panels A and B are from states +++ε and +++δ, respectively. (C) The PMF profiles from δ- and ε-nitrogen protonated His254 AWH simulations in +++ state. (D) and (E) Schematic drawing of different hydrogen bond patterns due to the δ- and ε-tautomeric states of His254.

### Energetics of the Histidine Tautomers

We also find that the tautomeric states of charge neutral histidine (δ- or ε-nitrogen protonated) can have a strong influence on His254 dynamics and energetics (Figs. 3, S2 and S3). For instance, in state +++δ, His254 orients to form a stable interaction with Ser150, whereas the other tautomer (ε-nitrogen protonated) is unstable in the B position (Fig. 3C). We rationalize this using the difference in extended hydrogen bonding patterns in the two states involving the ε-nitrogen of histidine, whereas the Ser150-His254 hydrogen bond via δ- nitrogen is likely hindered geometrically and sterically. In δ-nitrogen protonated state, His254 stabilizes in the B position with the ε-nitrogen acting as a hydrogen bond acceptor from Ser150, which in turn participates in a stable hydrogen bond network towards protonated Lys229 (Fig. 3E). However, when the ε-nitrogen of His254 is protonated, Ser150 does not act as hydrogen bond acceptor anymore, thereby leading to perturbation in the water-based hydrogen bonding with Lys229 and destabilization of the B conformation of histidine (Fig. 3D). Similarly, in other protonation states simulated, the histidine switch dynamics is found to be tautomer dependent. In 0+0 (or 000) charge state of lysine residues, His254 with ε-nitrogen protonated stabilizes in the B position by about 5 kcal/mol in comparison to the δ-nitrogen protonated state (see Figs. 2 and S2).

Previous simulation studies on Mrp antiporter showed that the doubly protonated histidine can occupy an intermediate position between the A and B conformations (Lee et al. 2022; Pecorilla et al. 2025). We modelled His254 in its imidazolium state and observed that it preferentially occupies the middle position, especially when Lys342 is protonated (+00), with both A and B positions at least 4 kcal/mol higher in energy (Fig. 4A). The effect of charge states of Lys305 and Lys229 on protonated histidine is relatively weak, and His254 occupies a position close to the A conformation, whereas the B position is unfavourable by at least 8 kcal/mol in all states studied (Fig. S3). A closer analysis of simulation trajectories reveals that a consecutive charge-assisted hydrogen bond network can be formed by protonated Lys342, Thr312 and neutral His254, explaining why the PMF-profile is stabilized in the A (or A-like) – conformation at an approx. RC ∼ -6 Å. In this scenario, His254 acts as a hydrogen-bond acceptor for the hydroxyl group of Thr312, whereas this is no longer possible in the imidazolium state of histidine. With His254 being positioned on a flexible α-helical segment, it therefore shifts away from the A conformation to an in-between position at an approx. RC ∼ -1.5 Å (Figs. 4B and 4C) or to similar values in other charge states (see Fig. S3).

**Figure 4.**
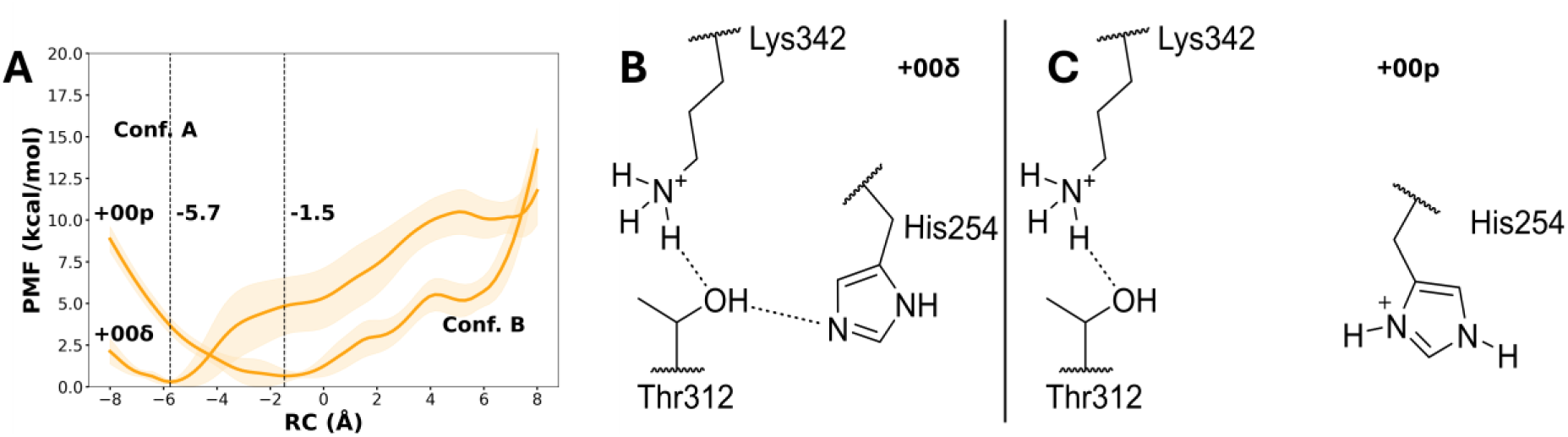
Protonation of neutral His254 causes a shift of the energy minimum within the PMF profile. (A) PMF profiles of histidine switch dynamics in two protonation states. The imidazolium state has its minimum shifted away from A position towards a state in between A and B. Panels (B) and (C) provide molecular and chemical explanations of the shift. With His254 in charge neutral state, it accepts a hydrogen bond from Thr312, which is no longer possible when histidine is doubly protonated. This, as well as the electrostatic repulsion from protonated Lys342, drives the protonated histidine towards the in-between states (see also Fig. S3).

The RCI histidine switch model (Pecorilla et al. 2025) suggests that His254 acts as a bridging element between the putative proton transfer paths (Fig. 1) and that it can minimize the formation of networks that leads to energy loss in terms of proton transfer down the gradient. Fig. 5 shows that in conformation A, the charge neutral His254 (Nδ or Nε tautomer) can bridge the solvent exposed Lys305 with the buried Lys342 via a hydrogen bonding network involving conserved Thr312 and Ser250, and water molecules. This pathway is reminiscent of a putative proton uptake route from the N side of the membrane. Since this His254-bridged connection is delinked from Lys229 of the Lys-Glu pair in the central hydrophilic axis, a likely function of the histidine switch is to prevent the premature protonation of anionic species generated upon quinone reduction at the other end of the complex.(Djurabekova et al. 2024) Furthermore, when His254 is in its B conformation, it participates in a path from protonated Lys229 (of Lys-Glu pair) to the δ-nitrogen of the histidine via two water molecules and Ser150 (Fig. 5). In this arrangement, the Lys342 and Lys305 remain disconnected, thereby effectively preventing a P-side to N-side connection.

**Figure 5.**
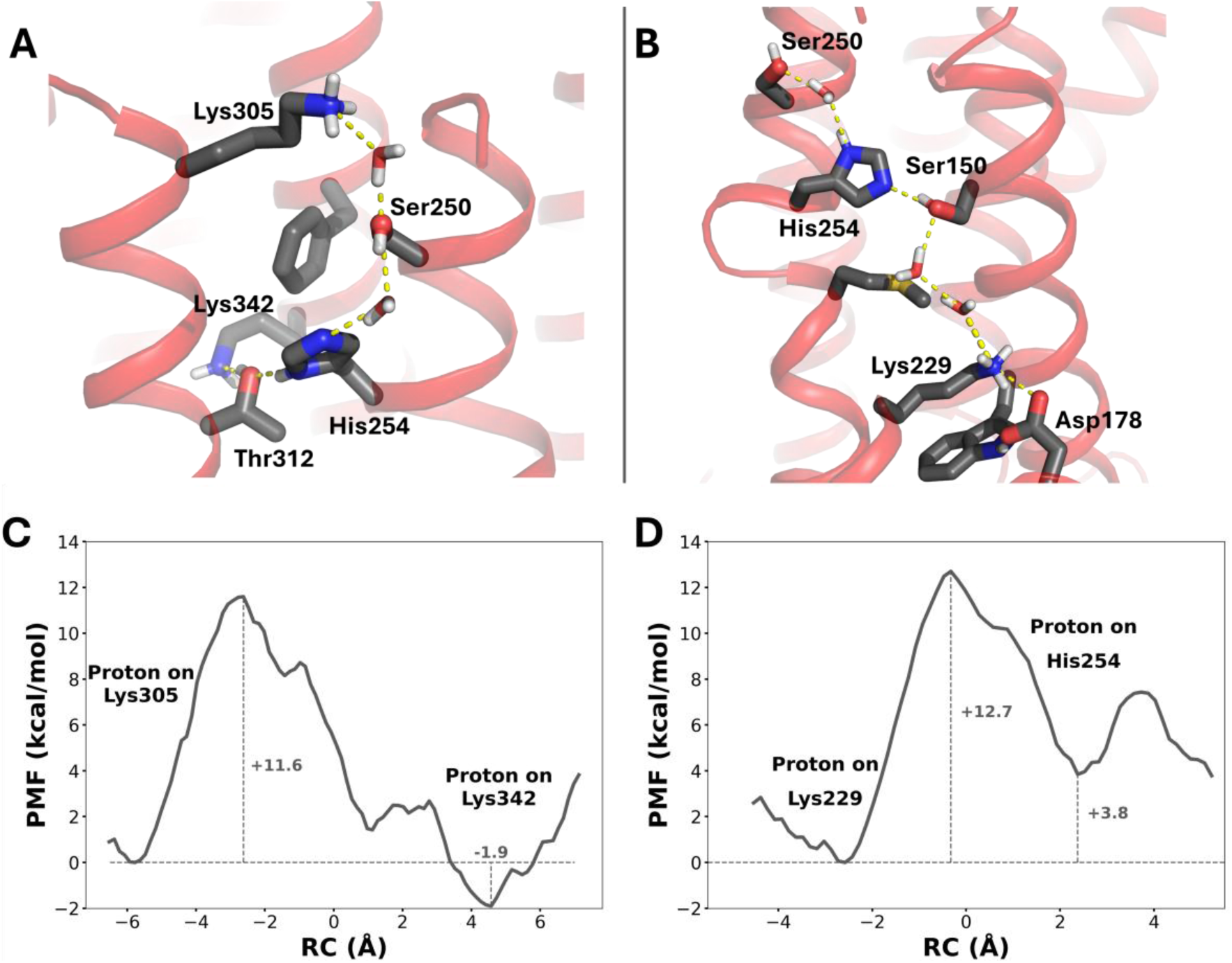
Simulation snapshots showcasing the hydrogen bonding patterns observed in simulation setups 0+0δ (A) and +++δ (B) with histidine in conformation A and B, respectively. The hydrogen bonds are shown with dotted lines. Parts of transmembrane helices are not shown for clarity. (C) PMF profile of proton transfer from Lys305 to Lys342 involving histidine in A position (panel A) and (D) from Lys299 to His254 in its B position (panel B) as a function of RC (in Å). See SI for description on RC.

### QM/MM free energy simulations of proton transfer

To investigate if the observed hydrogen bonded pathways (Fig. 5A and 5B) can catalyze proton transfer, we performed well-tempered metadynamics QM/MM free energy simulations (see Computational Methods). We observed that a proton can be transferred from the cytoplasm facing protonated Lys305 to the buried and charge neutral Lys342 involving histidine in A position (Fig. 5C). This proton transfer step may correspond to the loading of a pumped proton to the NuoL subunit from the cytoplasmic side in response to the quinone reduction reaction ca. 20 nm away. Several lines of evidence suggest a long-range electron-proton coupling in RCI and the mutation of Lys342 to alanine reduces the quinone reductase activity by ∼90%.(Sato et al. 2013) The histidine switch in A position effectively gates the transfer of cytoplasmic proton towards the quinone reduction site, which can cause uncoupling. When the proton transfer between histidine in B position and Lys229 is probed by QM/MM simulations, we find the protonation of His254 to be somewhat endergonic. It is indeed known from classical MD simulations that protonated histidine can stabilize a position in-between the two conformations A and B (Djurabekova, Haapanen, and Sharma 2020; Pecorilla et al. 2025), and such dynamical effects and long-range electrostatic effects are challenging to account for, which could bring the energetics favorable for histidine protonation. The activation energy barriers of ca. 12 kcal/mol, which roughly corresponding to 10-100 μs of proton transfer kinetics, do not compete with the complex I turnover (milliseconds). Our QM/MM calculations demonstrate that histidine can participate in proton transfer reactions and at the same time act as a molecular gate to prevent uncoupling.

## Discussion

All proposed models (Sazanov 2023; Djurabekova et al. 2024; Kaila 2018) suggest NuoL is involved in proton pumping by RCI. Being the most distant subunit from the quinone active site (Fig. 1), it must possess unique structural elements that enable efficient long-range coupling between the quinone reduction reaction and proton pumping. The highly conserved His254 is one such feature that may assist in efficient long-range coupling in RCI (Djurabekova, Haapanen, and Sharma 2020). Based on our enhanced sampling simulations and QM/MM approaches, we find that His254 undergoes protonation-state dependent conformational dynamics, shifts between putative proton transfer pathways (Fig. 1) and also acts as a proton transfer element. By stabilizing in the A conformation, His254 effectively shuts off the connectivity of the central hydrophilic axis with the proton transfer pathways of the terminal antiporter-like subunit of RCI, while in the B conformation, it detaches the proton uptake and release routes of the NuoL subunit. Furthermore, in the A position, it participates in proton uptake from the cytoplasmic side, whereas in B conformation, it can function as a proton carrier from the hydrophilic axis of RCI. Essentially, by switching between A and B conformations, His254 carries protons from the N side and the central hydrophilic axis to a proton release route towards the P side, thereby functioning as an efficient proton confurcation device. Overall, our multiscale simulation data strongly suggest that His254 is in fact a type of molecular switch that is capable of turning proton transfer routes on and off. The protonation states of the surrounding lysine residues (Lys229, Lys305 and Lys342) play a key role in triggering this switch, by rearranging water- and amino acid residue-based hydrogen bond networks. Further experimental testing by site-directed mutagenesis and cryo-EM studies will be required to strengthen the proposed model of histidine molecular switch.

The conservancy of this histidine extends beyond RCI and Mrp antiporter sequences, where it is also found in photosynthetic complex I (Zhang et al. 2020) and energy converting hydrogenase (Ech) (Katsyv and Müller 2022). Intriguingly, it is not conserved in this locus in membrane-bound hydrogenase (Mbh) (Yu et al. 2018), membrane-bound sulfane sulfur reductase (Mbs) (Yu et al. 2020) and formate hydrogenlyase (Fhl) enzymes (Steinhilper et al. 2022). However, the possibility of functional compensation by a histidine from another region cannot be slighted. Indeed, in agreement with this view the Fhl structure from *E. coli* shows a histidine residue (His222, *E. coli* Fhl numbering, PDB 7Z0S) that occupies a similar location and may take up the role of the histidine switch (Fig. S4). Further analysis of these putative switches in other protein classes is a subject of future studies.

### Computational methods

The 2.4 Å cryo-EM structure of *E. coli* RCI (PDB 7P7C) (Kravchuk et al. 2022) was used to construct the simulation model system comprising the five membrane-bound core subunits (NuoL, -M, -N, -K, and -J). A truncated model was prepared to keep the system size tractable and achieve enhanced simulation sampling in different chemical and conformational states (see below). Previously, truncated models of RCI have been simulated successfully providing functional insights (Galemou Yoga et al. 2020; Lasham et al. 2023; Nishida et al. 2025). The missing loops of NuoL and NuoN were added using Modeller (Šali and Blundell 1993). Protonation states were defined using PropKa (Søndergaard et al. 2011), with the exception of His254, Lys305, Lys342 and Lys229, which were simulated also in their alternative protonation states leading to in total of 24 different simulation setups (see Table S1). The OPM-aligned (Lomize et al. 2012) protein complex was embedded in a lipid bilayer composed of 75% POPE, 20% POPG and 5% dianionic cardiolipin, obtained using CHARMM-GUI (Jo et al. 2008). The membrane-protein system was solvated in a TIP3P water box with a salt concentration of 0.15 M Na^+^/Cl^-^. The CHARMM36 forcefield (Huang and MacKerell 2013) was used to describe the protein, lipids, ions and solvent. The model system comprised approx. 300 000 atoms (Fig. S5).

All classical atomistic MD simulations were performed with Gromacs-2022.4 (Abraham et al. 2015). Prior to production simulations, the system was energy minimized with a harmonic constraint of 2000 kJ mol^-1^ nm^-2^ on the protein heavy atoms. The resulting structure was equilibrated with a multi-step procedure, in analogy to prior literature (Lee et al. 2022): First, the temperature was equilibrated to 310 K for 100 ps in an NVT ensemble using the v-rescale thermostat (Bussi, Donadio, and Parrinello 2007). During this step, the constraints of the system were extended to the phosphorus atoms of lipids to prevent their movement in the z-direction. In the second step, the system was equilibrated in an NPT ensemble for 1 ns using the Berendsen barostat (Berendsen et al. 1984) (1 atm) and a v-rescale thermostat (Bussi, Donadio, and Parrinello 2007) (310 K) with the restraints from the phosphorus atoms removed. As a last equilibration step, all restraints were removed, and the system was simulated for 10 ns.

For the production simulations, the Parrinello-Rahman barostat (Parrinello and Rahman 1981) and the Nose-Hoover thermostat (Evans and Holian 1985) were employed to maintain a pressure of 1 atm and 310 K temperature, respectively. The long-range electrostatics was accounted for with PME (Darden, York, and Pedersen 1993) and time step of 2 fs was used using LINCS algorithm (Hess et al. 1997; Darden, York, and Pedersen 1993) implemented in Gromacs. A 12 Å cutoff for non-bonded interactions was used. Three 500 ns unbiased MD simulations were performed for each state to obtain starting coordinates for the AWH (Lindahl, Lidmar, and Hess 2014) enhanced sampling simulations with default parameters. The biased reaction coordinate for the AWH simulations is described in Fig. S6. 4 walkers were employed in each AWH run, and 4 replicas were simulated for each protonation state considered. The convergence of AWH simulation was achieved in ca. 200 ns (see Figs. S7). The total unbiased MD simulation sampling is 36 μs and AWH simulations corresponds to ca. 20 μs, totaling ∼56 μs of atomistic simulation sampling. The simulation data was analyzed using VMD (William, Andrew, and Klaus 1996), the Gromacs toolbox (Abraham et al. 2015) and MDAnalysis (Gowers et al. 2016; Michaud-Agrawal et al. 2011). Figures were produced using PyMol (Schrodinger 2015).

Snapshots for putative proton transfer pathways (Figs. 5A and 5B) were extracted from the AWH simulation trajectories to perform the well-tempered metadynamics-based QM/MM free energy calculations. The QM region consisted of the amino acid residues and water molecules displayed in Fig. 5 as well as the surroundings (see Table S3). For QM/MM simulations, NAMD (Phillips et al. 2020) and Orca (Neese et al. 2020) software were used. For the MM region, the CHARMM36 forcefield (Huang and MacKerell 2013) was employed, and the QM region was described by DFT with B3LYP (Becke 1993; Lee, Yang, and Parr 1988; Vosko, Wilk, and Nusair 1980; Stephens et al. 2002) functional def2-SVP (Weigend and Ahlrichs 2005) basis set. The dispersion effects were accounted for by using D3BJ (Grimme et al. 2010; Grimme, Ehrlich, and Goerigk 2011). The link-atom approach was used (link atom placed between CA and CB atom of the amino acid residues) together with the additive electrostatic embedding framework as implemented in NAMD. The QM/MM model system was energy minimized with 300 steps, followed by 1 ps of equilibration for both setups. Production runs were of 30 and 20 ps with well-tempered metadynamics for setups A and B, respectively (see Fig. 5). The hill height, width and ΔT were 0.3 kcal/mol, 0.3 Å and 4808 K, respectively. A cutoff of 12 Å was applied for non-bonded interactions with switching function at 10 Å. The temperature (310 K) was maintained with Langevin thermostat implemented in NAMD. The reaction coordinate to bias proton transfer was applied as discussed in Fig. S8. The *colvars* module (Fiorin, Klein, and Hénin 2013) was used to define the reaction coordinate. The metadynamics simulations were paused after the reaction coordinate sampled the defined path and the PMF profiles were stabilized (Fig. S9).

## Supplementary material

Supplementary Figures S1-S9 and Supplementary Tables S1-S3 can be found in the Supporting Information.

## Acknowledgements

VS acknowledges research funding from the Research Council of Finland, the Jane and Aatos Erkko Foundation, the Sigrid Jusélius Foundation, the University of Helsinki, the Cancer Foundation and the Finnish Society of Sciences and Letters. We acknowledge high-quality HPC resources from the Center for Scientific Computing (CSC), Finland. We are also thankful to the HiLIFE internship programme and to the IT support of the Faculty of Science, University of Helsinki. This work was supported by the German Research Foundation (Deutsche Forschungsgemeinschaft, CRC1507/P14 to VZ).

## References

1. Abraham, M.J., T. Murtola, R. Schulz, S. Páll, J.C. Smith, B. Hess and E. Lindahl. 2015. GROMACS: High performance molecular simulations through multi-level parallelism from laptops to supercomputers. SoftwareX 1–2:19–25.

2. Amarneh, B. and S.B. Vik. 2003. Mutagenesis of Subunit N of theEscherichia coliComplex I. Identification of the Initiation Codon and the Sensitivity of Mutants to Decylubiquinone. Biochemistry 42:4800–4808.

3. Becke, A.D. 1993. Density-functional thermochemistry. III. The role of exact exchange. The Journal of Chemical Physics 98:5648–5652.

4. Berendsen, H.J.C., J.P.M. Postma, W.F. van Gunsteren, A. DiNola and J.R. Haak. 1984. Molecular dynamics with coupling to an external bath. The Journal of Chemical Physics 81:3684–3690.

5. Bussi, G., D. Donadio and M. Parrinello. 2007. Canonical sampling through velocity rescaling. The Journal of Chemical Physics 126.

6. Darden, T., D. York and L. Pedersen. 1993. Particle mesh Ewald: An N⋅log(N) method for Ewald sums in large systems. The Journal of Chemical Physics 98:10089–10092.

7. Djurabekova, A., O. Haapanen and V. Sharma. 2020. Proton motive function of the terminal antiporter-like subunit in respiratory complex I. Biochimica et Biophysica Acta (BBA) - Bioenergetics 1861.

8. Djurabekova, A., J. Lasham, O. Zdorevskyi, V. Zickermann and V. Sharma. 2024. Long-range electron proton coupling in respiratory complex I — insights from molecular simulations of the quinone chamber and antiporter-like subunits. Biochemical Journal 481:499–514.

9. Euro, L., G. Belevich, M.I. Verkhovsky, M. Wikström and M. Verkhovskaya. 2008. Conserved lysine residues of the membrane subunit NuoM are involved in energy conversion by the proton-pumping NADH:ubiquinone oxidoreductase (Complex I). Biochimica et Biophysica Acta (BBA) - Bioenergetics 1777:1166–1172.

10. Evans, D.J. and B.L. Holian. 1985. The Nose–Hoover thermostat. The Journal of Chemical Physics 83:4069–4074.

11. Fiorin, G., M.L. Klein and J. Hénin. 2013. Using collective variables to drive molecular dynamics simulations. Molecular Physics 111:3345–3362.

12. Galemou Yoga, E., K. Parey, A. Djurabekova, O. Haapanen, K. Siegmund, K. Zwicker, V. Sharma, V. Zickermann and H. Angerer. 2020. Essential role of accessory subunit LYRM6 in the mechanism of mitochondrial complex I. Nature Communications 11.

13. Gowers, R., M. Linke, J. Barnoud, T. Reddy, M. Melo, S. Seyler, J. Domański, D. Dotson, S. Buchoux, I. Kenney and O. Beckstein. 2016. MDAnalysis: A Python Package for the Rapid Analysis of Molecular Dynamics Simulations. In Proceedings of the 15th Python in Science Conference, 98–105.

14. Grimme, S., J. Antony, S. Ehrlich and H. Krieg. 2010. A consistent and accurate ab initio parametrization of density functional dispersion correction (DFT-D) for the 94 elements H-Pu. The Journal of Chemical Physics 132.

15. Grimme, S., S. Ehrlich and L. Goerigk. 2011. Effect of the damping function in dispersion corrected density functional theory. Journal of Computational Chemistry 32:1456– 1465.

16. Hess, B., H. Bekker, H.J.C. Berendsen and J.G.E.M. Fraaije. 1997. LINCS: A linear constraint solver for molecular simulations. Journal of Computational Chemistry 18:1463–1472.

17. Hoeser, F., P. Saura, C. Harter, V.R.I. Kaila and T. Friedrich. 2025. A leigh syndrome mutation perturbs long-range energy coupling in respiratory complex I. Chemical Science 16:7374–7386.

18. Huang, J. and A.D. MacKerell. 2013. CHARMM36 all-atom additive protein force field: Validation based on comparison to NMR data. Journal of Computational Chemistry 34:2135–2145.

19. Jo, S., T. Kim, V.G. Iyer and W. Im. 2008. CHARMM-GUI: A web-based graphical user interface for CHARMM. Journal of Computational Chemistry 29:1859–1865.

20. Kaila, V.R.I. 2018. Long-range proton-coupled electron transfer in biological energy conversion: towards mechanistic understanding of respiratory complex I. Journal of The Royal Society Interface 15.

21. Kaila, V.R.I., M.I. Verkhovsky, G. Hummer and M. Wikström. 2008. Glutamic acid 242 is a valve in the proton pump of cytochrome c oxidase. Proceedings of the National Academy of Sciences 105:6255–6259.

22. Katsyv, A. and V. Müller. 2022. A purified energy-converting hydrogenase from Thermoanaerobacter kivui demonstrates coupled H+-translocation and reduction in vitro. Journal of Biological Chemistry 298.

23. Kravchuk, V., O. Petrova, D. Kampjut, A. Wojciechowska-Bason, Z. Breese and L. Sazanov. 2022. A universal coupling mechanism of respiratory complex I. Nature 609:808–814.

24. Lasham, J., O. Haapanen, V. Zickermann and V. Sharma. 2023. Tunnel dynamics of quinone derivatives and its coupling to protein conformational rearrangements in respiratory complex I. Biochimica et Biophysica Acta (BBA) - Bioenergetics 1864.

25. Lee, C., W. Yang and R.G. Parr. 1988. Development of the Colle-Salvetti correlation-energy formula into a functional of the electron density. Physical Review B 37:785–789.

26. Lee, Y., O. Haapanen, A. Altmeyer, W. Kühlbrandt, V. Sharma and V. Zickermann. 2022. Ion transfer mechanisms in Mrp-type antiporters from high resolution cryoEM and molecular dynamics simulations. Nature Communications 13.

27. Lindahl, V., J. Lidmar and B. Hess. 2014. Accelerated weight histogram method for exploring free energy landscapes. The Journal of Chemical Physics 141.

28. Lomize, M.A., I.D. Pogozheva, H. Joo, H.I. Mosberg and A.L. Lomize. 2012. OPM database and PPM web server: resources for positioning of proteins in membranes. Nucleic Acids Research 40:D370–D376.

29. Mathiesen, C. and C. Hägerhäll. 2002. Transmembrane topology of the NuoL, M and N subunits of NADH:quinone oxidoreductase and their homologues among membrane-bound hydrogenases and bona fide antiporters. Biochimica et Biophysica Acta (BBA) - Bioenergetics 1556:121–132.

30. Michaud-Agrawal, N., E.J. Denning, T.B. Woolf and O. Beckstein. 2011. MDAnalysis: A toolkit for the analysis of molecular dynamics simulations. Journal of Computational Chemistry 32:2319–2327.

31. Mitchell, P. 1961. Coupling of Phosphorylation to Electron and Hydrogen Transfer by a Chemi- Osmotic type of Mechanism. Nature 191:144–148.

32. Nakamaru-Ogiso, E., M.-C. Kao, H. Chen, S.C. Sinha, T. Yagi and T. Ohnishi. 2010. The Membrane Subunit NuoL(ND5) Is Involved in the Indirect Proton Pumping Mechanism of Escherichia coli Complex I. Journal of Biological Chemistry 285:39070–39078.

33. Neese, F., F. Wennmohs, U. Becker and C. Riplinger. 2020. The ORCA quantum chemistry program package. The Journal of Chemical Physics 152.

34. Nishida, M., C. Pecorilla, T. Masuya, K. Hirano, M. Abe, O. Zdorevskyi, V. Sharma, H. Miyoshi and M. Murai. 2025. Dynamic binding of acetogenin-type inhibitors to mitochondrial complex I revealed by photoaffinity labeling. Biochimica et Biophysica Acta (BBA) - Bioenergetics 1866.

35. Parey, K., C. Wirth, J. Vonck and V. Zickermann. 2020. Respiratory complex I — structure, mechanism and evolution. Current Opinion in Structural Biology 63:1–9.

36. Parrinello, M. and A. Rahman. 1981. Polymorphic transitions in single crystals: A new molecular dynamics method. Journal of Applied Physics 52:7182–7190.

37. Pecorilla, C., A. Altmeyer, O. Haapanen, Y. Lee, V. Zickermann and V. Sharma. 2025. Conformational dynamics of a histidine molecular switch in a cation/proton antiporter. Biochimica et Biophysica Acta (BBA) - Bioenergetics 1866.

38. Phillips, J.C., D.J. Hardy, J.D.C. Maia, J.E. Stone, J.V. Ribeiro, R.C. Bernardi, R. Buch, G. Fiorin, J. Hénin, W. Jiang, R. McGreevy, M.C.R. Melo, B.K. Radak, R.D. Skeel, A. Singharoy, Y. Wang, B. Roux, A. Aksimentiev, Z. Luthey-Schulten, L.V. Kalé, K. Schulten, C. Chipot and E. Tajkhorshid. 2020. Scalable molecular dynamics on CPU and GPU architectures with NAMD. The Journal of Chemical Physics 153.

39. Šali, A. and T.L. Blundell. 1993. Comparative Protein Modelling by Satisfaction of Spatial Restraints. Journal of Molecular Biology 234:779–815.

40. Sato, M., P.K. Sinha, J. Torres-Bacete, A. Matsuno-Yagi and T. Yagi. 2013. Energy Transducing Roles of Antiporter-like Subunits in *Escherichia coli* NDH-1 with Main Focus on Subunit NuoN (ND2). Journal of Biological Chemistry 288:24705–24716.

41. Sazanov, L.A. 2023. From the ‘black box’ to ‘domino effect’ mechanism: what have we learned from the structures of respiratory complex I. Biochemical Journal 480:319– 333.

42. Schrodinger, LLC. 2015. The PyMOL Molecular Graphics System, Version 1.8.

43. Søndergaard, C.R., M.H.M. Olsson, M. Rostkowski and J.H. Jensen. 2011. Improved Treatment of Ligands and Coupling Effects in Empirical Calculation and Rationalization of pKa Values. Journal of Chemical Theory and Computation 7:2284– 2295.

44. Steinhilper, R., G. Höff, J. Heider and B.J. Murphy. 2022. Structure of the membrane-bound formate hydrogenlyase complex from Escherichia coli. Nature Communications 13.

45. Stephens, P.J., F.J. Devlin, C.F. Chabalowski and M.J. Frisch. 2002. Ab Initio Calculation of Vibrational Absorption and Circular Dichroism Spectra Using Density Functional Force Fields. The Journal of Physical Chemistry 98:11623–11627.

46. Swartz, T.H., S. Ikewada, O. Ishikawa, M. Ito and T.A. Krulwich. 2005. The Mrp system: a giant among monovalent cation/proton antiporters? Extremophiles 9:345–354.

47. Verkhovskaya, M.L., N. Belevich, L. Euro, M. Wikström and M.I. Verkhovsky. 2008. Real-time electron transfer in respiratory complex I. Proceedings of the National Academy of Sciences 105:3763–3767.

48. Vosko, S.H., L. Wilk and M. Nusair. 1980. Accurate spin-dependent electron liquid correlation energies for local spin density calculations: a critical analysis. Canadian Journal of Physics 58:1200–1211.

49. Weigend, F. and R. Ahlrichs. 2005. Balanced basis sets of split valence, triple zeta valence and quadruple zeta valence quality for H to Rn: Design and assessment of accuracy. Physical Chemistry Chemical Physics 7.

50. Wikström, M., C. Pecorilla and V. Sharma. 2023. The mitochondrial respiratory chain. In *History of The Enzymes*, Current Topics and Future Perspectives, 15–36.

51. William, H., D. Andrew and S. Klaus. 1996. VMD -- V isual M olecular D ynamics. Journal of Molecular Graphics 14:33–38.

52. Yang, S. and Q. Cui. 2011. Glu-286 Rotation and Water Wire Reorientation Are Unlikely the Gating Elements for Proton Pumping in Cytochrome c Oxidase. Biophysical Journal 101:61–69.

53. Yu, H., D.K. Haja, G.J. Schut, C.-H. Wu, X. Meng, G. Zhao, H. Li and M.W.W. Adams. 2020. Structure of the respiratory MBS complex reveals iron-sulfur cluster catalyzed sulfane sulfur reduction in ancient life. Nature Communications 11.

54. Yu, H., C.-H. Wu, G.J. Schut, D.K. Haja, G. Zhao, J.W. Peters, M.W.W. Adams and H. Li. 2018. Structure of an Ancient Respiratory System. Cell 173:1636–1649.e1616.

55. Zhang, C., J. Shuai, Z. Ran, J. Zhao, Z. Wu, R. Liao, J. Wu, W. Ma and M. Lei. 2020. Structural insights into NDH-1 mediated cyclic electron transfer. Nature Communications 11.

