## Supporting Information for "Molecular and Energetic Basis of Histidine Switch Dynamics in Respiratory Complex I"

**Fig. S1. Scatter plot showing the position of the histidine in various 3D structures of complex I and related proteins within the OPM data bank**. Proteins that could not be structurally aligned with the membrane-embedded domain of *E. coli* complex I (PDB 7P7C) or those that did not have histidine present or resolved (17 of 177) were excluded from the plot. See Table S2 for full list of proteins considered and Figure S6 for a depiction of the considered distances.

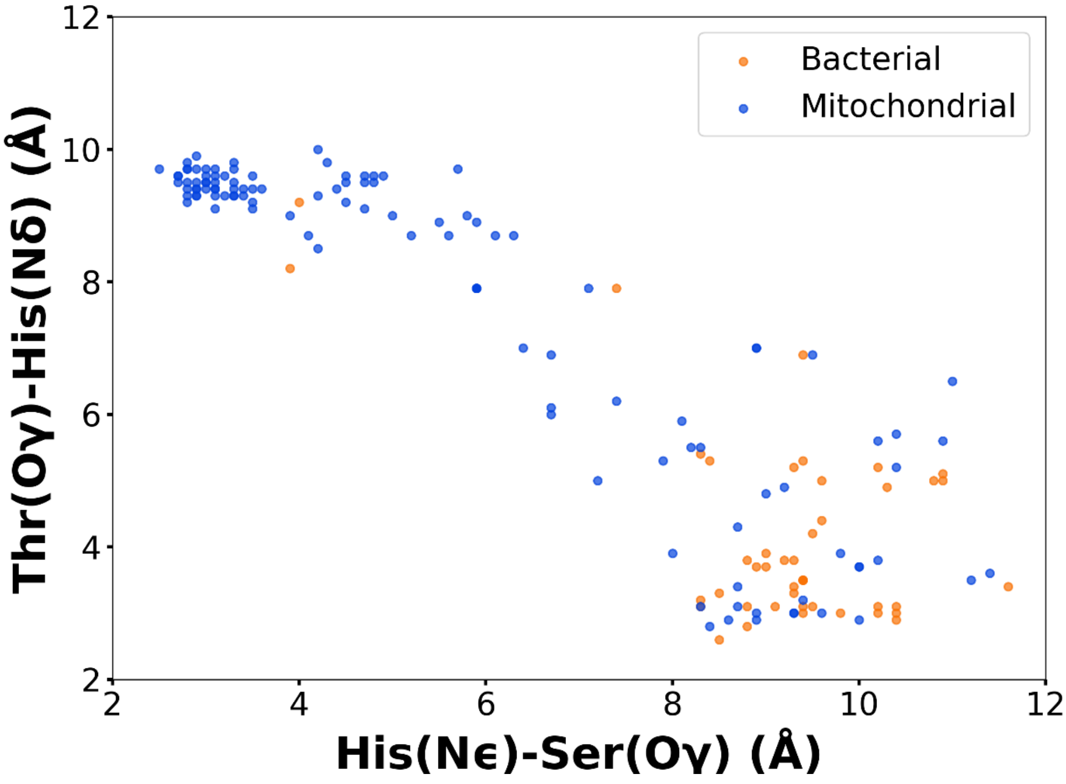

**Fig. S2. Energetics of charge-neutral ε-nitrogen protonated histidine sidechain dynamics.** The y-axis in panels (A) and (B) describe the potential of mean force (PMF, kcal/mol) with respect to the reaction coordinate (RC) on x-axis in Å (see methods). Each trace is the mean of four simulation replicas, whereas the shaded region describes the standard error of mean. See also Fig. 1 in main text. Notations for the protonation states are described in Fig. 2 of the main text.

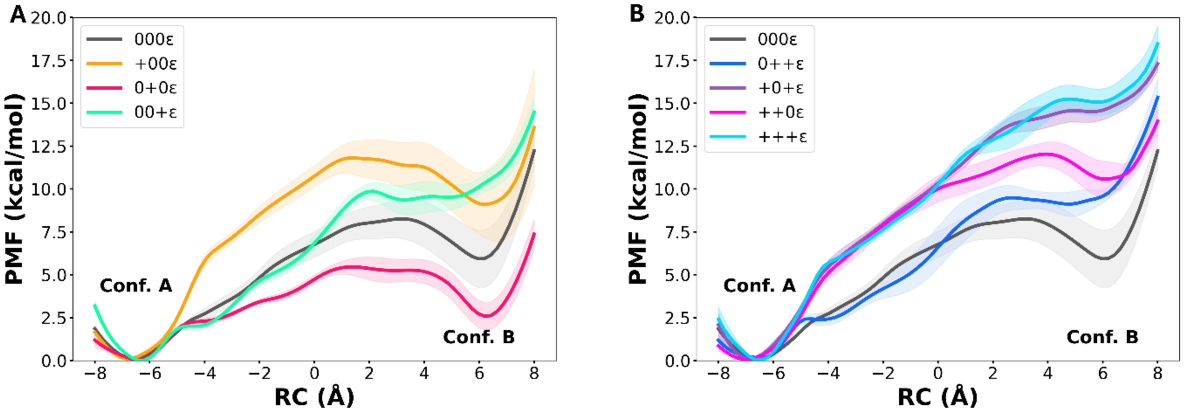

**Fig. S3. Energetics of doubly protonated His254 sidechain conformational dynamics.** The y-axis in panels (A) and (B) describe the potential of mean force (PMF, kcal/mol) with respect to the reaction coordinate (RC) on x-axis in Å (see methods). Each trace is the mean of four simulation replicas, whereas the shaded region describes the standard error of mean. See also Fig. 1 in main text.

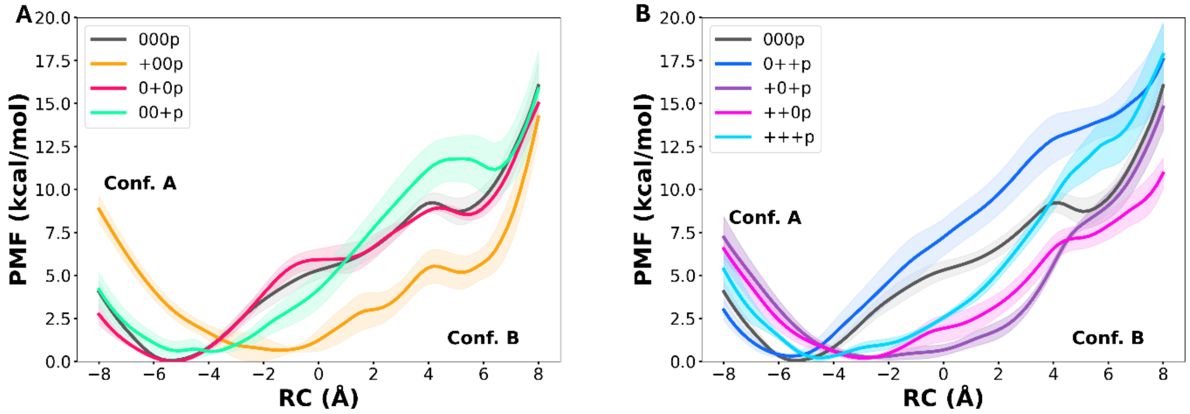

**Fig. S4. Histidine switch substitution in complex I like protein families.** Residues in *E. coli* complex I (PDB 7P7C, red cartoon and grey residues) and membrane-bound Fhl (7Z0S, orange cartoon and cyan residues) occupying similar spatial locations. The membrane-bound Fhl possesses Ser234 instead of the conserved His254, however, His222 from a neighboring transmembrane helix occupies a similar position as the putative histidine switch. With Lys342/Lys336 and Thr312/Thr292 being conserved as well, Fhl shows the same residues as the putative proton transfer pathway in RCI.

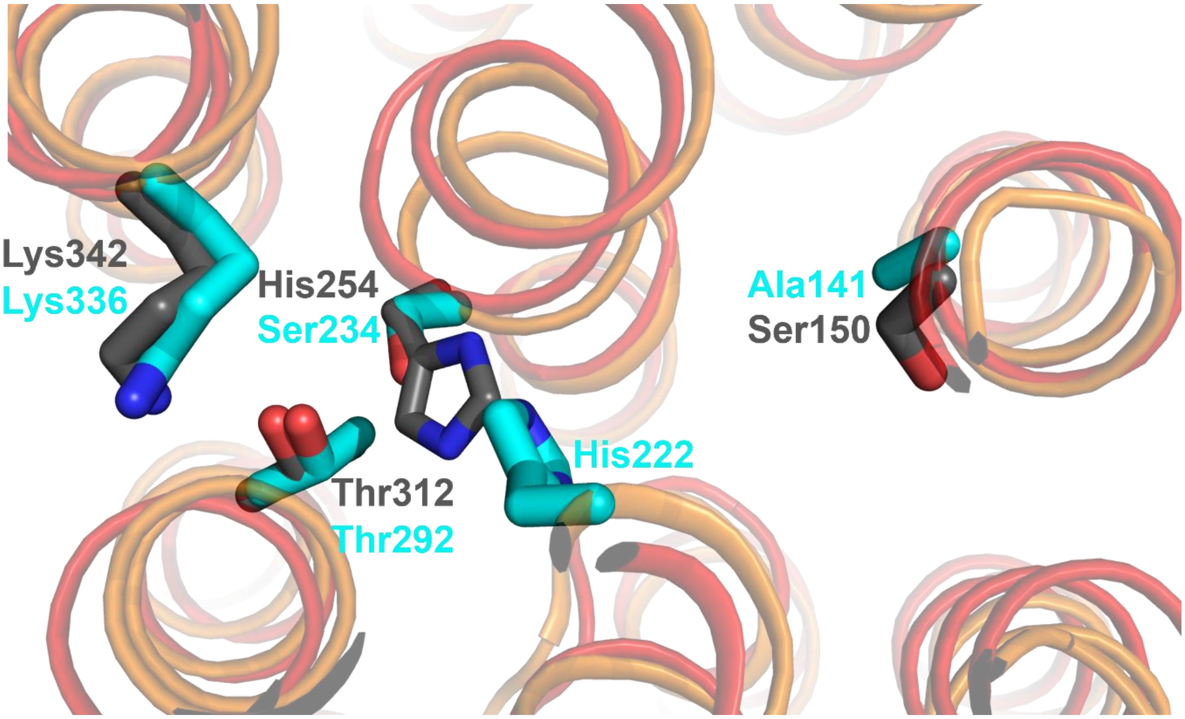

**Fig. S5. The protein model immersed in lipid bilayer.** In this top view from cytoplasmic side, the protein subunits are shown in colored ribbons. The lipids (shades of green, see methods) surrounding the protein are shown with sticks. The $Na^+$ and $Cl^-$ ions are displayed as blue and orange spheres, respectively. The water solvent is omitted for clarity.

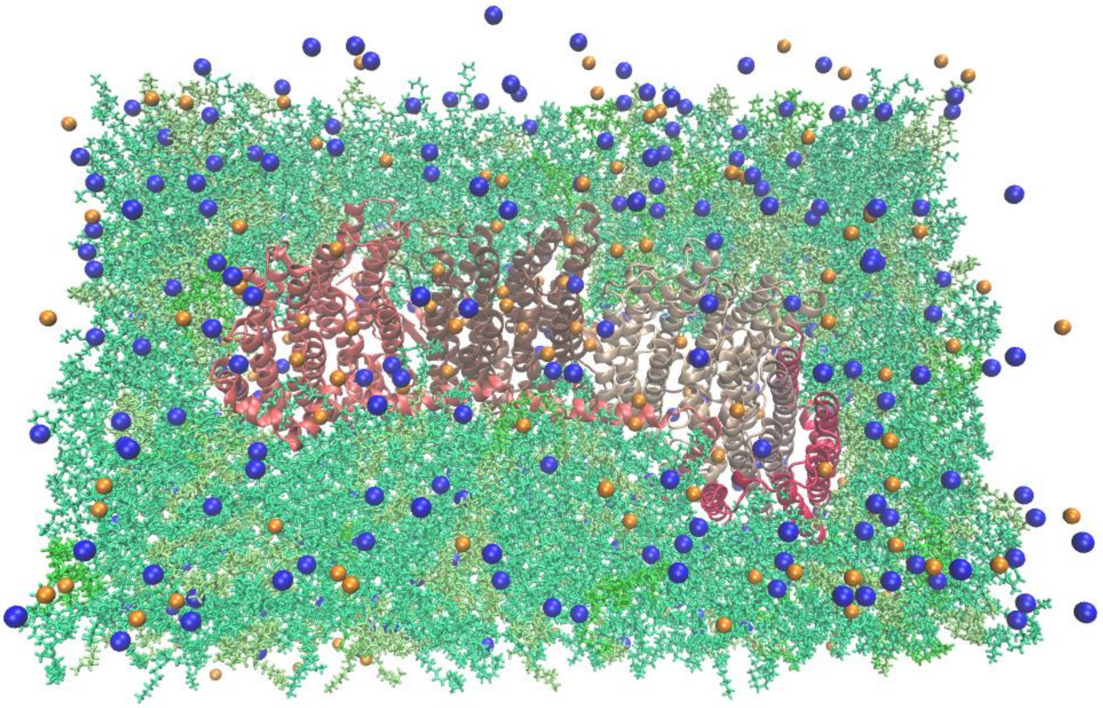

**Fig. S6. Reaction coordinate used in AWH simulations.** Conformation A is characterized by a hydrogen bond between the Thr312-Hydroxyl oxygen atom and the $N_\delta$-atom of His254 ($r_A$), while Conformation B is defined by a hydrogen bond between the Ser150-hydroxyl oxygen atom and the $N_\varepsilon$-atom of His254 ($r_B$). The reaction coordinate was then defined as RC = $r_A - r_B$, so that negative values correspond to being closer to conformation A, while positive values correspond to being closer to conformation B. As a guideline, we defined RC <= -0.6 nm as conformation A (e.g. PDB 7P7C shows an RC of -0.58 nm), while RC >= 0.6 nm corresponds to conformation B (e.g. PDB 7QRU shows an RC of 0.63 nm). The sampling interval was defined from -0.8 nm to 0.8 nm. It is noteworthy that no forces are directly applied to $r_A$ or $r_B$, thus, the imidazole moiety of His254 can rotate freely during the transition from A to B.

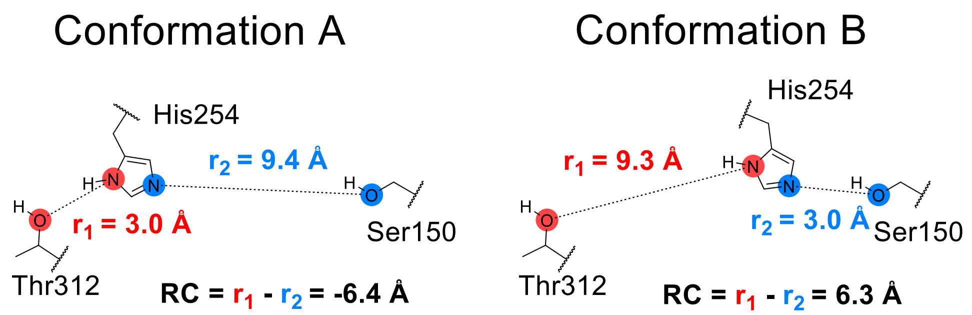

**Fig. S7. Convergence of AWH simulations.** The top panels show behaviour of four independent AWH simulation replicas for two selected protonation states. Across 24 different simulation setups there are many cases in which the variation in simulation replicas is rather small. We selected these to point out that variation across replicas can be seen in a simulation setup, but overall behaviour of PMF profile is consistent across replicas. The lower panels show time dependency of the PMF profile for selected replicas of two simulation setups, highlighting convergence.

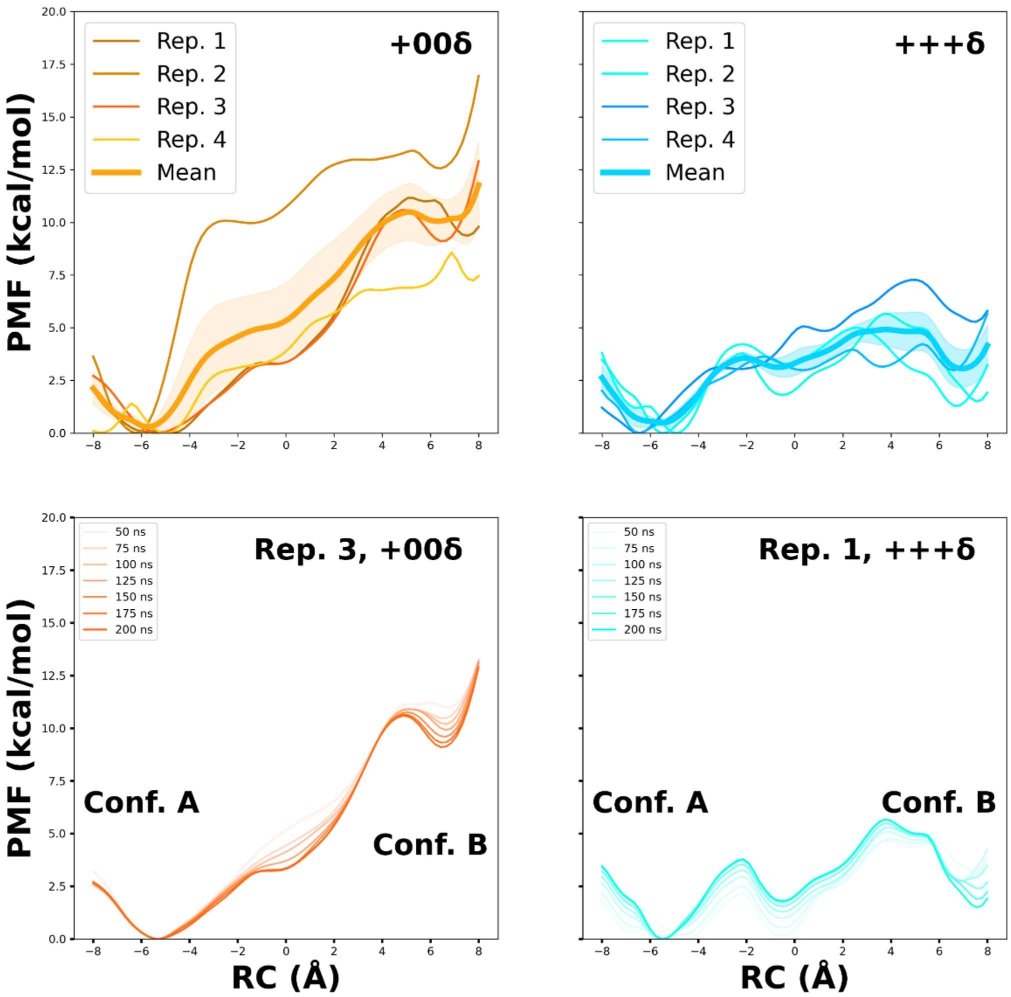

**Fig. S8. Reaction coordinate used in QM/MM simulations.** For simulating proton transfer from donor (protonated lysine sidechain) to acceptor (neutral lysine sidechain) via water molecules and polar residues, a reaction coordinate (RC) was sampled with metadynamics approach. RC corresponds to summing the differences of bond distances of hydrogen from donor and acceptor for every hydrogen bond in the pathway. RC = (sum of *red* distances) minus (sum of *blue* distances).

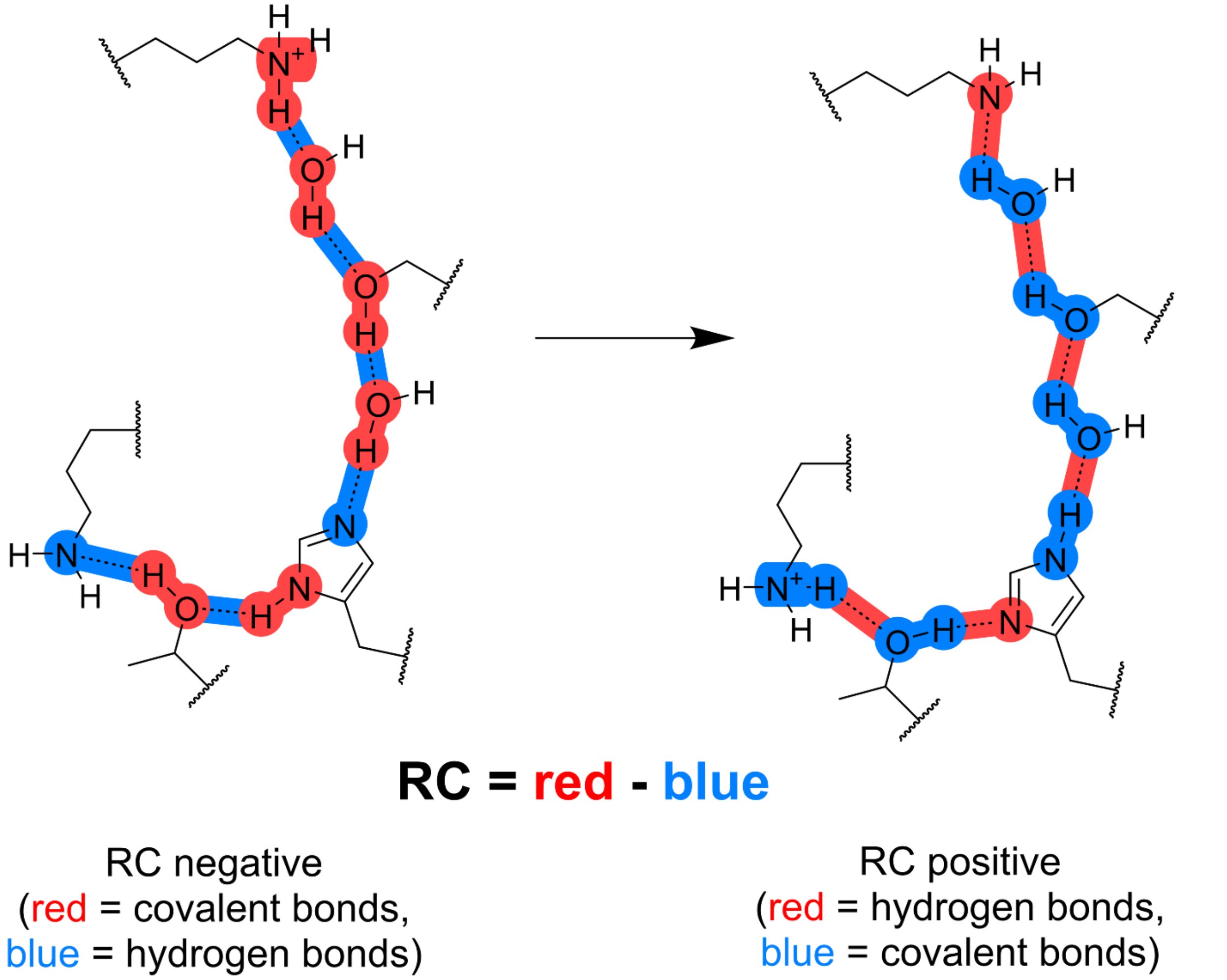

**Fig. S9. Convergence of well-tempered metadynamics-based QM/MM free energy simulations.** Top panels display the sampling of the reaction coordinate in well-tempered metadynamics for simulations based on snapshot A and B. Lower panels show convergence of PMF profile with respect to simulation time.

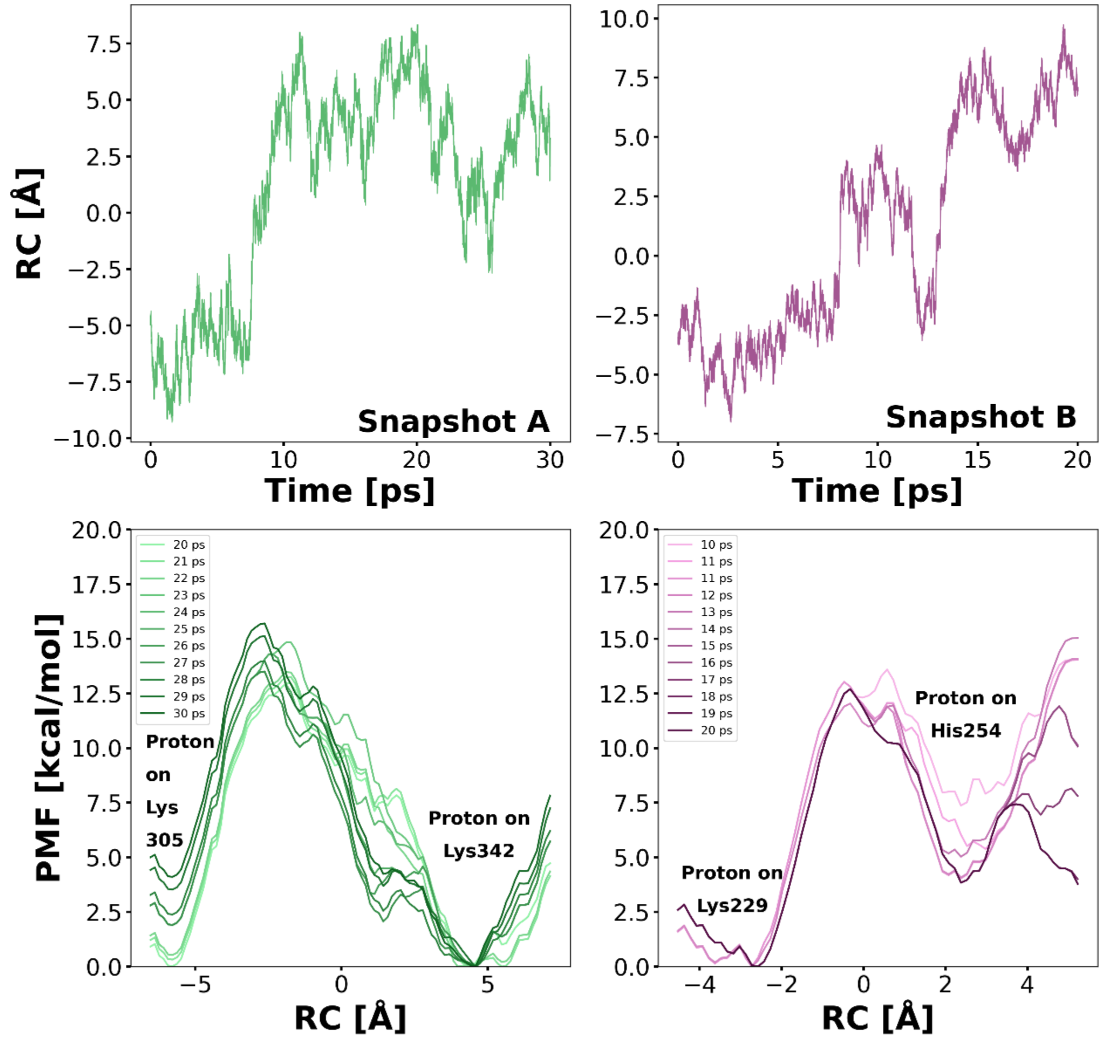

**Table S1. Model systems for classical MD simulations.** The table summarizes all combinations of protonation states investigated in this work. All setups were simulated for 3 x 500 ns in an unbiased manner and with AWH for 4 x 200 ns.

| Lys342 | Lys305 | Lys229 | His254 |
|---|---|---|---|
| 0 | + | + | δ |
| 0 | + | + | ε |
| 0 | + | + | p |
| 0 | 0 | + | δ |
| 0 | 0 | + | ε |
| 0 | 0 | + | p |
| + | + | + | δ |
| + | + | + | ε |
| + | + | + | p |
| 0 | + | 0 | δ |
| 0 | + | 0 | ε |
| 0 | + | 0 | p |
| 0 | 0 | 0 | δ |
| 0 | 0 | 0 | ε |
| 0 | 0 | 0 | p |
| + | + | 0 | δ |
| + | + | 0 | ε |
| + | + | 0 | p |
| + | 0 | 0 | δ |
| + | 0 | 0 | ε |
| + | 0 | 0 | p |
| + | 0 | + | δ |
| + | 0 | + | ε |
| + | 0 | + | p |

**Table S2. Location of histidine in X-ray and cryo-EM structures of RCI and related proteins.**

| PDB | Resolution (Å) | Organism | Thr-His(ND1) (Å) | Ser-His(NE2) (Å) | RC (Å) |
|---|---|---|---|---|---|
| 7b93 | 3.04 | *Mus musculus* | 9.7 | 5.7 | 4 |
| 7b0n | 3.7 | *Yarrowia lipolytica* | 9.4 | 4.4 | 5 |
| 7v2e | 2.8 | *Sus scrofa* | 9.5 | 4.8 | 4.7 |
| 7ard | 3.11 | *Polytomella sp.* | 3.8 | 10.2 | -6.4 |
| 7qsl | 2.76 | *Bos taurus* | 9.6 | 2.7 | 6.9 |
| 7qsk | 2.84 | *Bos taurus* | 9.7 | 2.5 | 7.2 |
| 8b9z | 3.28 | *Drosophila melanogaster* | 6.9 | 9.5 | -2.6 |
| 7v2h | 2.5 | *Sus scrofa* | 9.5 | 2.8 | 6.7 |
| 7v2c | 2.9 | *Sus scrofa* | 9.7 | 3 | 6.7 |
| 7v2r | 2.6 | *Sus scrofa* | 9.5 | 4.5 | 5 |
| 7v31 | 2.9 | *Sus scrofa* | 9.5 | 2.9 | 6.6 |
| 7v33 | 2.6 | *Sus scrofa* | 9.8 | 3.3 | 6.5 |
| 7z7r | 3.36 | *Escherichia coli* | 2.6 | 8.5 | -5.9 |
| 7p7c | 2.4 | *Escherichia coli* | 5.1 | 10.9 | -5.8 |
| 7p62 | 3.6 | *Escherichia coli* | 4.9 | 10.3 | -5.4 |
| 7p7e | 2.7 | *Escherichia coli* | 2.9 | 10.4 | -7.5 |
| 7zdj | 3.25 | *Ovis aries* | 8.9 | 5.5 | 3.4 |
| 7zeb | 3.8 | *Ovis aries* | 9.9 | 2.9 | 7 |
| 7zdp | 3.88 | *Ovis aries* | 9.3 | 4.2 | 5.1 |
| 7qso | 3.02 | *Bos taurus* | 9.2 | 3.5 | 5.7 |
| 7qsd | 3.1 | *Bos taurus* | 3.9 | 8 | -4.1 |
| 7qsn | 2.81 | *Bos taurus* | 9.4 | 2.9 | 6.5 |
| 7qsm | 2.3 | *Bos taurus* | 9.4 | 2.9 | 6.5 |
| 7v3m | 2.9 | *Sus scrofa* | 9.4 | 2.8 | 6.6 |
| 7v2k | 2.7 | *Sus scrofa* | 9.5 | 4.7 | 4.8 |
| 7v2d | 3.3 | *Sus scrofa* | 9.6 | 4.9 | 4.7 |
| 7v2f | 3.1 | *Sus scrofa* | 10 | 4.2 | 5.8 |
| 7v30 | 2.7 | *Sus scrofa* | 9.6 | 2.7 | 6.9 |
| 7v32 | 3.2 | *Sus scrofa* | 9.6 | 4.8 | 4.8 |
| 6zkc | 3.1 | *Ovis aries* | 9.3 | 3.1 | 6.2 |
| 6zkd | 2.7 | *Ovis aries* | 9.6 | 3.2 | 6.4 |
| 6zke | 2.6 | *Ovis aries* | 9.8 | 2.8 | 7 |
| 6zkf | 2.8 | *Ovis aries* | 9.7 | 2.9 | 6.8 |
| 7zkq | 3.15 | *Yarrowia lipolytica* | complex I assembly intermediate | | |
| 8esw | 3.3 | *Drosophila melanogaster* | 5.7 | 10.4 | -4.7 |
| 7zdm | 3.44 | *Ovis aries* | 9 | 5 | 4 |
| 7zdh | 3.46 | *Ovis aries* | 9.6 | 3.5 | 6.1 |
| 7zd6 | 3.16 | *Ovis aries* | 9.2 | 4.5 | 4.7 |
| 8esz | 3.4 | *Drosophila melanogaster* | 5.3 | 7.9 | -2.6 |

| | | | | | |
|---|---|---|---|---|---|
| 7p7m | 3.2 | *Escherichia coli* | 5.2 | 9.3 | -4.1 |
| 6zkk | 3.7 | *Ovis aries* | 9.4 | 3.3 | 6.1 |
| 6zkl | 3.1 | *Ovis aries* | 9.3 | 3.2 | 6.1 |
| 6zkm | 2.8 | *Ovis aries* | 9.7 | 2.8 | 6.9 |
| 6zkn | 2.9 | *Ovis aries* | 9.7 | 2.8 | 6.9 |
| 7zkp | 3.2 | *Yarrowia lipolytica* | complex I assembly intermediate | | |
| 7ar9 | 2.97 | *Polytomella sp.* | 5.6 | 10.9 | -5.3 |
| 7vbl | 2.6 | *Sus scrofa* | 9.3 | 3.3 | 6 |
| 7vc0 | 2.6 | *Sus scrofa* | 9.4 | 3 | 6.4 |
| 7vwl | 2.7 | *Sus scrofa* | 9.5 | 2.7 | 6.8 |
| 7vbp | 2.8 | *Sus scrofa* | 9.6 | 4.7 | 4.9 |
| 8e9h | 2.7 | *Mycolicibacterium smegmatis* | 3.1 | 10.2 | -7.1 |
| 8e9i | 2.8 | *Mycolicibacterium smegmatis* | 5.2 | 10.2 | -5 |
| 8e9g | 2.6 | *Mycolicibacterium smegmatis* | 3 | 10.2 | -7.2 |
| 7o6y | 3.4 | *Yarrowia lipolytica* | 4.8 | 9 | -4.2 |
| 8ba0 | 3.68 | *Drosophila melanogaster* | 5.6 | 10.2 | -4.6 |
| 7z7s | 2.4 | *Escherichia coli* | 3.1 | 8.8 | -5.7 |
| 7p63 | 3.4 | *Escherichia coli* | 3.4 | 11.6 | -8.2 |
| 7z7v | 2.29 | *Escherichia coli* | 2.8 | 8.8 | -6 |
| 7z7t | 3.1 | *Escherichia coli* | 3 | 10.4 | -7.4 |
| 7p64 | 2.5 | *Escherichia coli* | 3.1 | 8.3 | -5.2 |
| 7zci | 2.69 | *Escherichia coli* | 3.1 | 9.5 | -6.4 |
| 7p69 | 3 | *Escherichia coli* | 3.1 | 9.1 | -6 |
| 7z80 | 2.93 | *Escherichia coli* | 3.8 | 8.8 | -5 |
| 7z84 | 2.87 | *Escherichia coli* | 3.2 | 8.3 | -5.1 |
| 7z83 | 2.88 | *Escherichia coli* | 3.3 | 8.5 | -5.2 |
| 7zc5 | 3 | *Escherichia coli* | 3.1 | 9.4 | -6.3 |
| 7p7j | 2.7 | *Escherichia coli* | 5.3 | 9.4 | -4.1 |
| 7p7k | 3.1 | *Escherichia coli* | 5.4 | 8.3 | -2.9 |
| 7p7l | 3 | *Escherichia coli* | 5.3 | 8.4 | -3.1 |
| 6zkg | 3.4 | *Ovis aries* | 9.4 | 3.1 | 6.3 |
| 6zkh | 3 | *Ovis aries* | 9.4 | 3.1 | 6.3 |
| 6zki | 2.8 | *Ovis aries* | 9.6 | 3.1 | 6.5 |
| 6zkj | 3 | *Ovis aries* | 9.4 | 3.6 | 5.8 |
| 7p61 | 3.2 | *Escherichia coli* | 5 | 10.8 | -5.8 |
| 7ak5 | 3.17 | *Mus Musculus* | 8.7 | 6.3 | 2.4 |
| 6zks | 3.1 | *Ovis aries* | 9.7 | 3.3 | 6.4 |
| 6zkt | 2.8 | *Ovis aries* | 9.5 | 3.1 | 6.4 |
| 6zku | 3 | *Ovis aries* | 9.3 | 3.4 | 5.9 |
| 6zkv | 2.9 | *Ovis aries* | 9.4 | 3.5 | 5.9 |
| 7z0t | 3.4 | *Escherichia coli* | Fhl, no histidine | | |
| 7z0s | 2.6 | *Escherichia coli* | Fhl, no histidine | | |
| 3i9v | 3.1 | *Thermus thermophilus* | complex I peripheral arm | | |

| | | | | | |
|---|---|---|---|---|---|
| 8bq6 | 2.8 | *Arabidopsis thaliana* | 9.1 | 3.1 | 6 |
| 6zkb | 2.9 | *Ovis aries* | 9.6 | 3 | 6.6 |
| 6zka | 2.5 | *Ovis aries* | 9.7 | 3.1 | 6.6 |
| 6x89 | 3.9 | *Vigna radiata* | His carrying subunit absent | | |
| 7zm7 | 2.77 | *Chaetomium thermophilum* | 9.8 | 4.3 | 5.5 |
| 7zm8 | 2.76 | *Chaetomium thermophilum* | 9.6 | 4.5 | 5.1 |
| 7zmg | 2.44 | *Chaetomium thermophilum* | 9.3 | 2.8 | 6.5 |
| 7zmh | 2.47 | *Chaetomium thermophilum* | 9.2 | 2.8 | 6.4 |
| 7zmb | 2.75 | *Chaetomium thermophilum* | 9 | 3.9 | 5.1 |
| 7zme | 2.83 | *Chaetomium thermophilum* | 9.1 | 4.7 | 4.4 |
| 5xth | 3.9 | *Homo sapiens* | 7.9 | 5.9 | 2 |
| 5gup | 4 | *Sus scrofa* | 8.5 | 4.2 | 4.3 |
| 5xtd | 3.7 | *Homo sapiens* | 7.9 | 5.9 | 2 |
| 5xti | 17.4 | *Homo sapiens* | 7.9 | 5.9 | 2 |
| 6z16 | 2.98 | *Anoxybacillus flavithermus* | 7.9 | 7.4 | 0.5 |
| 7qru | 2.24 | *Alkalihalobacillus pseudofirmus* | 2.8 or 9.3 | 8.8 or 3.0 | -6 or 6.3 |
| 7d3u | 3 | *Dietzia sp.* | 5 | 9.6 | -4.6 |
| 6zko | 3.8 | *Ovis aries* | 9.5 | 3 | 6.5 |
| 6zkp | 3.2 | *Ovis aries* | 9.5 | 3.3 | 6.2 |
| 6zkq | 3.3 | *Ovis aries* | 9.1 | 3.5 | 5.6 |
| 6zkr | 3.5 | *Ovis aries* | 9.4 | 3.4 | 6 |
| 7eu3 | 3.7 | *Hordeum vulgare* | 6.5 | 11 | -4.5 |
| 6zk9 | 2.3 | *Ovis aries* | complex I peripheral arm | | |
| 8bq5 | 2.73 | *Arabidopsis thaliana* | 9.5 | 3 | 6.5 |
| 8bef | 2.13 | *Arabidopsis thaliana* | His carrying subunit absent | | |
| 8bpx | 2.09 | *Brassica oleracea* | 9.3 | 2.9 | 6.4 |
| 8beh | 2.29 | *Brassica oleracea* | 9.3 | 2.9 | 6.4 |
| 7a23 | 3.7 | *Brassica oleracea* | 5.2 | 10.4 | -5.2 |
| 7a24 | 3.8 | *Brassica oleracea* | complex I assembly intermediate | | |
| 7ar8 | 3.53 | *Brassica oleracea* | 7.9 | 7.1 | 0.8 |
| 7arb | 3.41 | *Brassica oleracea* | 8.7 | 5.2 | 3.5 |
| 7aqq | 3.06 | *Brassica oleracea* | His carrying subunit absent | | |
| 7aqw | 3.17 | *Brassica oleracea* | 8.7 | 6.1 | 2.6 |
| 7ar7 | 3.72 | *Brassica oleracea* | 9 | 5.8 | 3.2 |
| 7nyh | 3.6 | *Escherichia coli* | 3.5 | 9.4 | -5.9 |
| 2ybb | 19 | *Bos taurus* | Sidechain not modeled | | |
| 7nyr | 3.3 | *Escherichia coli* | 3.5 | 9.4 | -5.9 |
| 6cfw | 3.7 | *Pyrococcus furiosus* | Mbh, histidine likely not conserved | | |
| 3rko | 3 | *Escherichia coli* | 5 | 10.9 | -5.9 |
| 7ak6 | 3.82 | *Mus musculus* | 8.7 | 4.1 | 4.6 |
| 7nyv | 3.7 | *Escherichia coli* | 3.5 | 9.4 | -5.9 |
| 7nyu | 3.8 | *Escherichia coli* | 3.5 | 9.4 | -5.9 |
| 6zr2 | 3.1 | *Mus musculus* | 8.7 | 5.6 | 3.1 |

| | | | | | |
|---|---|---|---|---|---|
| 6u8y | 4 | *Pyrococcus furiosus* | Mbs, histidine likely not conserved | | |
| 5ldw | 4.27 | *Bos taurus* | 3.7 | 10 | -6.3 |
| 5lc5 | 4.35 | *Bos taurus* | 3.9 | 9.8 | -5.9 |
| 5ldx | 5.6 | *Bos taurus* | 3.7 | 10 | -6.3 |
| 7dgz | 3.8 | *Bos taurus* | 6.1 | 6.7 | -0.6 |
| 6ztq | 3 | *Mus musculus* | 8.9 | 5.9 | 3 |
| 5o31 | 4.13 | *Bos taurus* | 4.3 | 8.7 | -4.4 |
| 6hum | 3.34 | *Thermosynechococcus elongatus* | 6.9 | 9.4 | -2.5 |
| 6nby | 3.1 | *Thermosynechococcus elongatus* | 3.9 | 9 | -5.1 |
| 6nbq | 3.1 | *Thermosynechococcus elongatus* | Sidechain not resolved | | |
| 6nbx | 3.5 | *Thermosynechococcus elongatus* | 3.8 | 9.2 | -5.4 |
| 6khi | 3 | *Thermosynechococcus elongatus* | 8.2 | 3.9 | 4.3 |
| 6khj | 3 | *Thermosynechococcus elongatus* | 9.2 | 4 | 5.2 |
| 6l7o | 3.2 | *Thermosynechococcus elongatus* | 4.2 | 9.5 | -5.3 |
| 6l7p | 3.6 | *Thermosynechococcus elongatus* | 4.4 | 9.6 | -5.2 |
| 6tjv | 3.2 | *Thermosynechococcus elongatus* | Arginine instead of histidine | | |
| 4wz7 | 3.6 | *Yarrowia lipolytica* | 3.5 | 11.2 | -7.7 |
| 6rfr | 3.2 | *Yarrowia lipolytica* | 5.5 | 8.3 | -2.8 |
| 6h8k | 3.79 | *Yarrowia lipolytica* | 3.6 | 11.4 | -7.8 |
| 6rfq | 3.3 | *Yarrowia lipolytica* | 5.5 | 8.2 | -2.7 |
| 6yj4 | 2.7 | *Yarrowia lipolytica* | 9.3 | 3.3 | 6 |
| 7dh0 | 4.2 | *Bos taurus* | 7 | 8.9 | -1.9 |
| 5lnk | 3.9 | *Ovis aries* | 4.9 | 9.2 | -4.3 |
| 4hea | 3.3 | *Thermus thermophilus* | 3 | 9.4 | -6.4 |
| 6y11 | 3.11 | *Thermus thermophilus* | 3.7 | 8.9 | -5.2 |
| 6i1p | 3.21 | *Thermus thermophilus* | 3.3 | 9.3 | -6 |
| 6i0d | 3.6 | *Thermus thermophilus* | 3 | 9.8 | -6.8 |
| 6q8o | 3.61 | *Thermus thermophilus* | 3.7 | 9 | -5.3 |
| 6q8w | 3.4 | *Thermus thermophilus* | 3.4 | 9.3 | -5.9 |
| 6q8x | 3.51 | *Thermus thermophilus* | 3.8 | 9.3 | -5.5 |
| 6ziy | 4.25 | *Thermus thermophilus* | 3.1 | 10.4 | -7.3 |
| 6zjy | 5.5 | *Thermus thermophilus* | Sidechain not resolved | | |
| 5xtc | 3.7 | *Homo sapiens* | 7.9 | 5.9 | 2 |
| 6qc6 | 4.1 | *Ovis aries* | 6.2 | 7.4 | -1.2 |
| 7r41 | 2.3 | *Bos taurus* | 3.1 | 8.3 | -5.2 |
| 7r42 | 2.3 | *Bos taurus* | 2.8 | 8.4 | -5.6 |

| | | | | | |
|---|---|---|---|---|---|
| 7r43 | 2.4 | *Bos taurus* | 3.1 | 8.7 | -5.6 |
| 7r44 | 2.4 | *Bos taurus* | 2.9 | 8.6 | -5.7 |
| 7r45 | 2.4 | *Bos taurus* | 3 | 9.6 | -6.6 |
| 7r46 | 2.4 | *Bos taurus* | 3.4 | 8.7 | -5.3 |
| 7r47 | 2.3 | *Bos taurus* | 3 | 8.9 | -5.9 |
| 7r48 | 2.3 | *Bos taurus* | 3.2 | 9.4 | -6.2 |
| 7r4c | 2.3 | *Bos taurus* | 3 | 9.3 | -6.3 |
| 7r4d | 2.3 | *Bos taurus* | 2.9 | 10 | -7.1 |
| 7r4f | 2.4 | *Bos taurus* | 3 | 9.3 | -6.3 |
| 7r4g | 2.5 | *Bos taurus* | 2.9 | 8.9 | -6 |
| 6q9b | 3.9 | *Ovis aries* | 7 | 6.4 | 0.6 |
| 6qa9 | 4.1 | *Ovis aries* | 6.9 | 6.7 | 0.2 |
| 6qc4 | 4.6 | *Ovis aries* | 5 | 7.2 | -2.2 |
| 7dgr | 4.6 | *Bos taurus* | 5.9 | 8.1 | -2.2 |
| 7dgs | 7.8 | *Bos taurus* | 6 | 6.7 | -0.7 |
| 7dkf | 8.3 | *Bos taurus* | 7 | 8.9 | -1.9 |

Table S3. QM/MM setups simulated in this work. Residues marked in red were part of the RC sampled.

| Snapshot | Protonation state | Number of Atoms | Residues in QM-Region | Total charge |
|---|---|---|---|---|
| A | 0+0δ | 168 | Leu239, Met243, Ser250, Ile253, His254(δ), Lys305(+), Ser311, Thr312, Lys342(0), His338(δ), Phe346, Glu359(0), 10+2 water molecules | +1 |
| B | +++δ | 172 | Trp143, Glu144(0), Ser150, Ile154, Phe171, Thr174, Arg175(+), Asp178(0), Lys229(+), Trp238, Ser250, His254(δ), Met258, 4+2 water molecules | +2 |
